# Significant functional differences despite morphological and molecular similarity in fully differentiated matched Conditionally Reprogrammed (CRC) and Feeder free dual SMAD inhibited expanded human nasal epithelial cells

**DOI:** 10.1101/2020.05.29.120006

**Authors:** Nikhil T. Awatade, Sharon L. Wong, Elvis Pandzic, Iveta Slapetova, Alexander Capraro, Ling Zhong, Nihan Turgutoglu, Laura K. Fawcett, Renee M. Whan, Adam Jaffe, Shafagh A. Waters

## Abstract

**Background:** Patient-derived airway cells differentiated at Air Liquid Interface (ALI) are valuable models for Cystic fibrosis (CF) precision therapy. Advances in culture techniques have improved expansion capacity of airway basal cells, while retaining functional airway epithelium physiology. However, considerable variation in response to CFTR modulators is observed even when using similar ALI culture techniques. We aimed to address if variation in response reflects true biological differences between patients or technical differences as a consequence of different culture expansion methods.

**Methods:** Nasal epithelial brushings from 14 individuals (CF=9; non-CF=5) were collected, then equally divided and expanded under conditional reprogramming culture (CRC) and feeder-serum-free “dual-SMAD inhibition” (SMADi) methods. Expanded cells from each culture were differentiated with proprietary PneumaCult™-ALI media. Morphology (Immunofluorescence), global proteomics (LC-MS/MS) and function (barrier integrity, cilia motility, and ion transport) were compared in CRC^ALI^ and SMADi^ALI^ under basal and CFTR corrector treated (VX-809) conditions.

**Results:** No significant difference in the structural morphology or global proteomics profile were observed. Barrier integrity and cilia motility were significantly different, despite no difference in cell junction morphology or cilia abundance. Epithelial Sodium Channels and Calcium-activated Chloride Channel activity did not differ but CFTR mediated chloride currents were significantly reduced in SMADi^ALI^ compare to their CRC^ALI^ counterparts.

**Conclusion:** Alteration of cellular physiological function *in vitro* occurs were more prominent than structural and differentiation potential in airway ALI. Since culture conditions significantly influence CFTR activity, this could lead to false conclusions if data from different labs are compared against each other without specific reference ranges.

## 1. INTRODUCTION

Following years of end-organ symptom treatment, novel targeted CFTR therapies (known as CFTR modulators) that can restore CFTR protein function have been developed. The large number (>2000) of variants in the CFTR gene results in a variety of clinical phenotypes and multiple different CFTR structural defects (1). CFTR modulators such as correctors that stabilize and increase CFTR protein trafficking (e.g. VX-809/Lumacaftor) and potentiators, which increase channel opening probability (e.g. VX-770/Ivacaftor) have gained regulatory approval to treat people with CF, with common and well-characterised CFTR mutations (2). Yet, inter-subject variability among individuals with the same CFTR genotype has been described. Patients with F508del, the most common CFTR mutation amongst the CF population worldwide, display a spectrum of responses to CFTR-modulator drugs despite having the same CFTR mutation variant (2, 3).

One of the key goals of the CF field has been the advancement of personalised therapies. Differentiated primary human bronchial epithelial cells (HBECs) have been instrumental for understanding CFTR structure and for testing rare CFTR variant- and patient-specific responsiveness to modulator drugs (4-7). HBECs grown at an air-liquid interface (ALI) is the gold standard pre-clinical model system for CF translational studies (8). Since HBECs can only be isolated through invasive procedures, sampling a large number of patients is challenging. Human nasal epithelial cells (HNECs) are increasingly shown to be an appropriate, non-invasive surrogate for HBECs since they share profound similarities in CFTR expression profile, growth characteristics and mucociliary differentiation pattern (9-12).

Different *in vitro* culture methods have been developed to overcome the limitation of primary cell proliferative capacity. Amongst these, the conditional reprogramming culture (CRC) technique is currently most widely used (13-15). CRC supports the long-term expansion of airway epithelial cells with the use of irradiated feeder cells and RhoA kinase inhibitor. More recently, a feeder and serum-free approach has been described for long-term clonal growth of HBEC. This approach uses small molecule inhibitors of the SMAD-dependent TGF-β (Transforming Growth Factor Beta) and Bone Morphogenic Protein (BMP) signalling pathways, referred to as “dual-SMAD inhibition” (SMADi) (16, 17). SMADi cultures confer the advantage of having no contaminating feeder cells. Both methods have been shown to enhance cell growth and lifespan while preserving electrophysiological and morphological properties (11, 16, 18-21).

A lack of well-defined standardised culture conditions for patient derived airway cells has led to considerable variation in cell differentiation observed between research groups when using similar ALI culture techniques. In studies where cells are sourced from various commercial vendors or academic biobanks, it is thus difficult to decipher if this reflects true biological differences between subjects or technical differences due to the manner or circumstances in which the cells were obtained and expanded. The objective of this study was to ascertain if using different methodology (CRC vs. SMADi) to expand primary nasal epithelial cells resulted in differences in molecular, structural and functional profile of those cells when differentiation at Air Liquid Interface (ALI) (**Fig 1A**).

**Fig 1.**
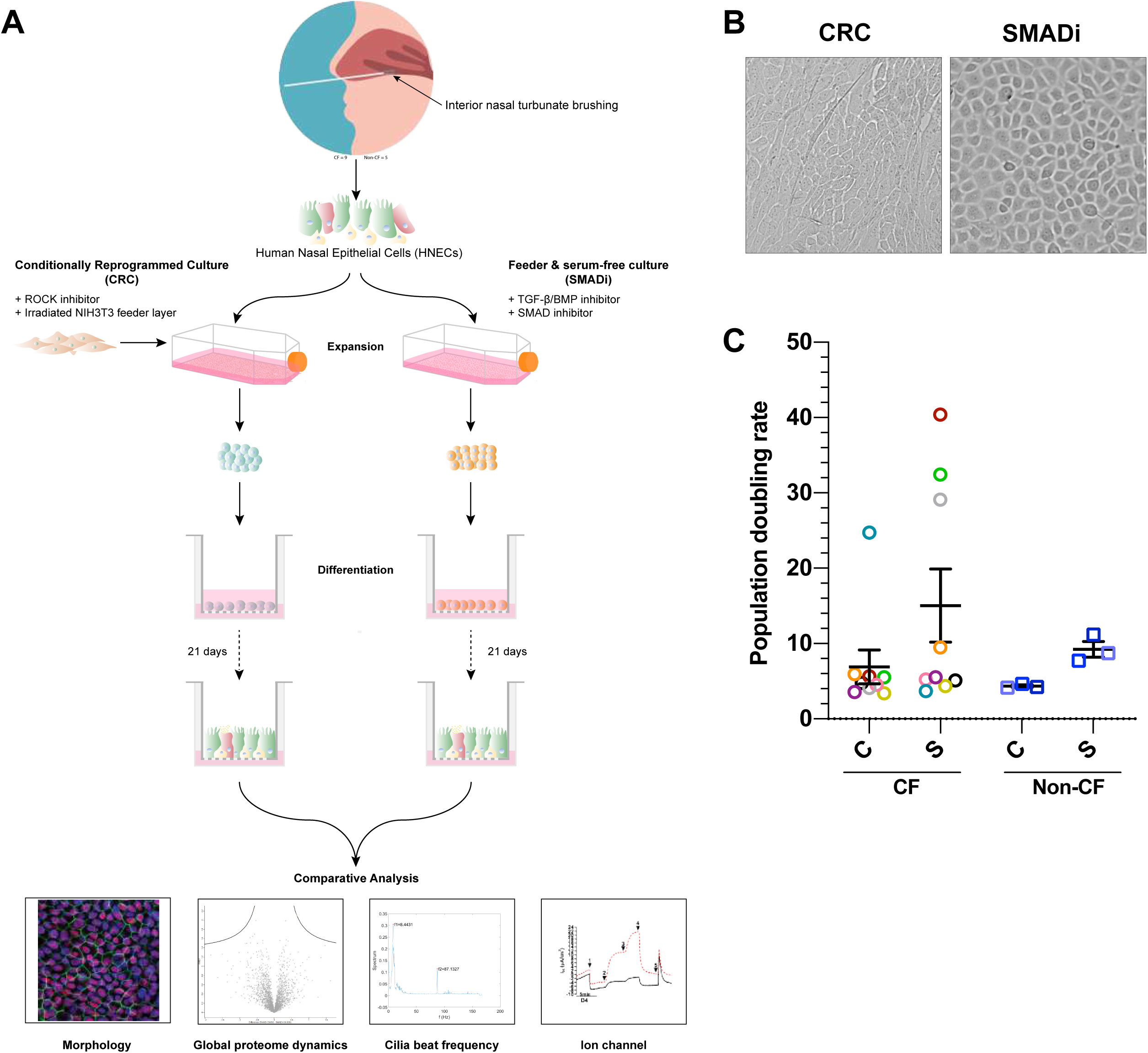
Expansion of human nasal epithelial (HNE) cells using conditional reprogramming culture (CRC) and dual SMAD inhibition (SMADi) method. **(A)** Study design schematic. **(B)** Representative image of CRC HNE cells cultured from a F508del/F508del CF patient using the CRC (left) and SMADi methods. Scale bars = 50 µm. Images are from Donor 5. **(C)** Population doubling rate of CF (n=9) and non-CF (n=5) HNE cells at passage 0. Each coloured circle represents an individual donor. Error bars represent standard error of the mean (Mean ± SEM). A two-tailed Student’s t-test was used to determine statistical significance. * = P ≤ 0.05.

In this study, we collected nasal epithelial brushings from 14 individuals (nine with CF and five non-CF controls). The brushed cells were divided and expanded with two distinct culture techniques. The CRC- and SMADi - expanded basal epithelial cells from each donor were then differentiated into mature pseudostratified epithelial cells at air-liquid interface with proprietary Pneumacult-ALI media. We found that despite no significant difference in the differentiated epithelium’s structural morphology or global proteomics profile between the two culture systems, their functional behaviour (assessed by epithelial barrier integrity, cilia beat frequency and ion channel activity) was significantly different.

## 2. MATERIALS and METHODS

Detailed materials and methods are available as supplementary materials (**Supplementary 1**).

## 3. RESULTS

Paired CRC- and SMADi-expanded HNE cells were created from 14 donors. Of these nine have homozygous F508del-CFTR, and five donors have wild-type CFTR genotype. SMADi cultures exhibited a neatly packed cobblestone morphology, characteristic of epithelial cells. CRC cultures demonstrated cells of similar size but the cells appeared to have a more oval shape (**Fig 1B**). Both CRC and SMADi cultures demonstrated donor-to-donor variability in two growth characteristics. On average, cultures reached ∼80% confluency in ∼7 to 20 days. The population doubling rate of CRC cultures were largely similar to those of the SMADi cultures, in both CF and non-CF cultures, averaging between 4.35 and 15.04 (**Fig 1C**). Both cultures demonstrated donor-to-donor variability in rate of growth, with a few notable slower-growing cultures in both methods.

### 3.1 Morphology of CRC and SMADi expanded cultures differentiated at air liquid interface is similar

Cells from each expansion method were cultured on porous membrane transwells with identical differentiation methodology. These cultures, hereafter referred to as CRC^ALI^ and SMADi^ALI^, both showed formation of a polarised, pseudostratified epithelial layer with mucociliary differentiation. No difference in differentiation potential was observed between donor paired CRC^ALI^ and SMADi^ALI^ cultures. Cells were uniform, organized, and had typical epithelial cobblestone morphology, with little variability observed in all ALI cultures (**Fig 2-** Donor 1; and **Fig S1-** Donor 4).

**Fig 2.**
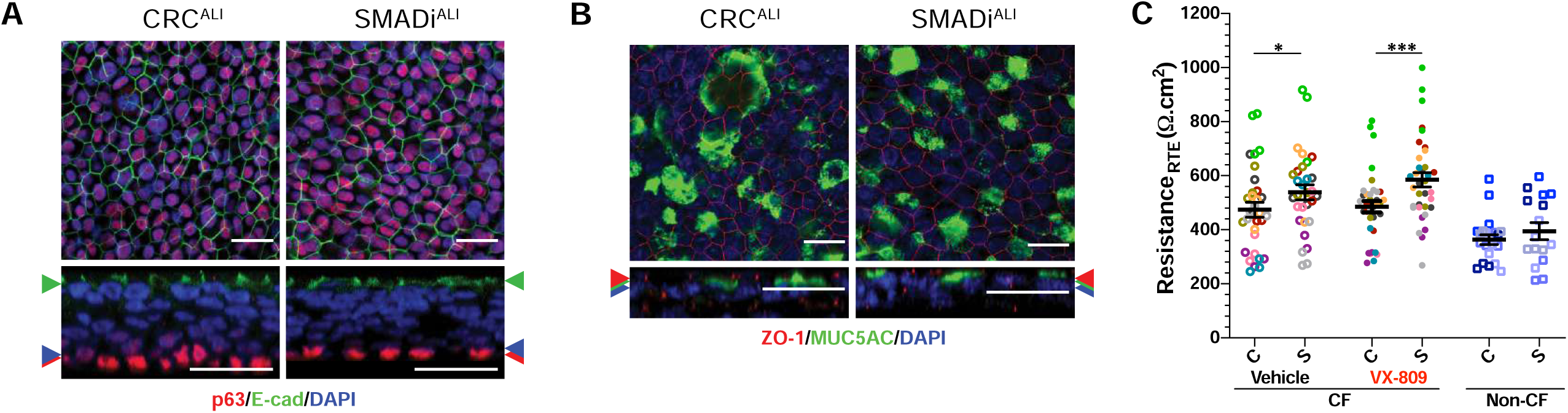
Structural characterisation of donor matched CRC and SMADi HNE cells grown at Air-Liquid Interface (ALI). Immunofluorescence staining of **(A)** basal progenitor cells p63+ (red) and adherens junctions, E-cadherin (green). **(B)** Mucus-producing goblet cells, MUC5AC (green) and tight junctions, ZO-1 (red). **(C)** Trans-epithelial electrical resistance (RTE) values of F508del/F508del CFTR (n=9) with and without VX-809 treat-ment and WT CFTR (n=5). XY-images shown in all panels are merged from single channel images acquired at Z-planes indicated by coloured arrows. 63x/1.4 oil immersion objective. Scale bars = 20µm. C = CRC, S = SMADi. Error bars represent standard error of the mean (Mean ± SEM). A two-tailed Student’s t-test was used to determine statistical significance. * = P ≤ 0.05 and *** = P ≤ 0.001.

To test the epithelial architecture, we performed an immunofluorescence characterisation of two CF donor-matched CRC^ALI^ and SMADi^ALI^ cultures (**Fig 2** and **Fig S1**). In CF CRC^ALI^ and SMADi^ALI^ cultures, stem cell p63 marker lined the basal cell compartment of the stratified epithelium (red; **Fig 2A**). Distribution of adherens junction, E-cadherin (green; **Fig 2A and Fig S1**) proteins and tight junction proteins, ZO-1 (red; **Fig 2B, Fig S1**) did not differ and was limited to the apical cells. These proteins are characteristic features of mature differentiated airway epithelia and demonstrate an intact epithelial barrier. Furthermore, we compared the trans-epithelial electrical resistance (R_TE_) of cultures as a quantitative technique to measure the integrity of tight junction dynamics in both ALI models. Both CRC^ALI^ and SMADi^ALI^ cultures exhibited donor-to-donor variability in resistance values, which ranged between 200 to 1000 Ω.cm^2^ (**Fig 2C, Fig S3A**). Irrespective of the CFTR genotype, a trend of higher resistance was observed in SMADi^ALI^ when compared to CRC^ALI^ cultures. In CF ALI cultures, this trend reached statistical significance (R_TE_: 538.7 ± 27.55 vs. 474.2 ± 27.55 Ω.cm^2^ respectively, P ≤ 0.01).

### 3.2 Global proteomic signature of CRC and SMADi expanded cultures differentiated at air liquid interface are similar

Assessing molecular differences between the CRC^ALI^ and SMADi^ALI^ cultures was achieved by performing global label-free proteomics analyses. CF and non-CF untreated cultures and CF cultures treated with VX-809 were assessed (**Fig 3**). Between CF and non-CF, 2305 and 2314 proteins were identified in CRC^ALI^ and SMADi^ALI^ cultures, respectively. There were no global differences in any proteins (**Fig 3)** including those specifically involved in SMADi or TGF-β regulation (**Table S4**) between CF and non-CF under either CRC^ALI^ or SMADi^ALI^ methods. In CF, between untreated CRC^ALI^ and SMADi^ALI^ cultures, 2505 proteins were identified, while between VX-809-treated CRC^ALI^ and SMADi^ALI^ cultures, 2514 proteins were identified. No differentially abundant proteins (q-value < 0.05, fold-change > 2) could be determined (**Fig 3 and Table S4**).

**Fig 3.**
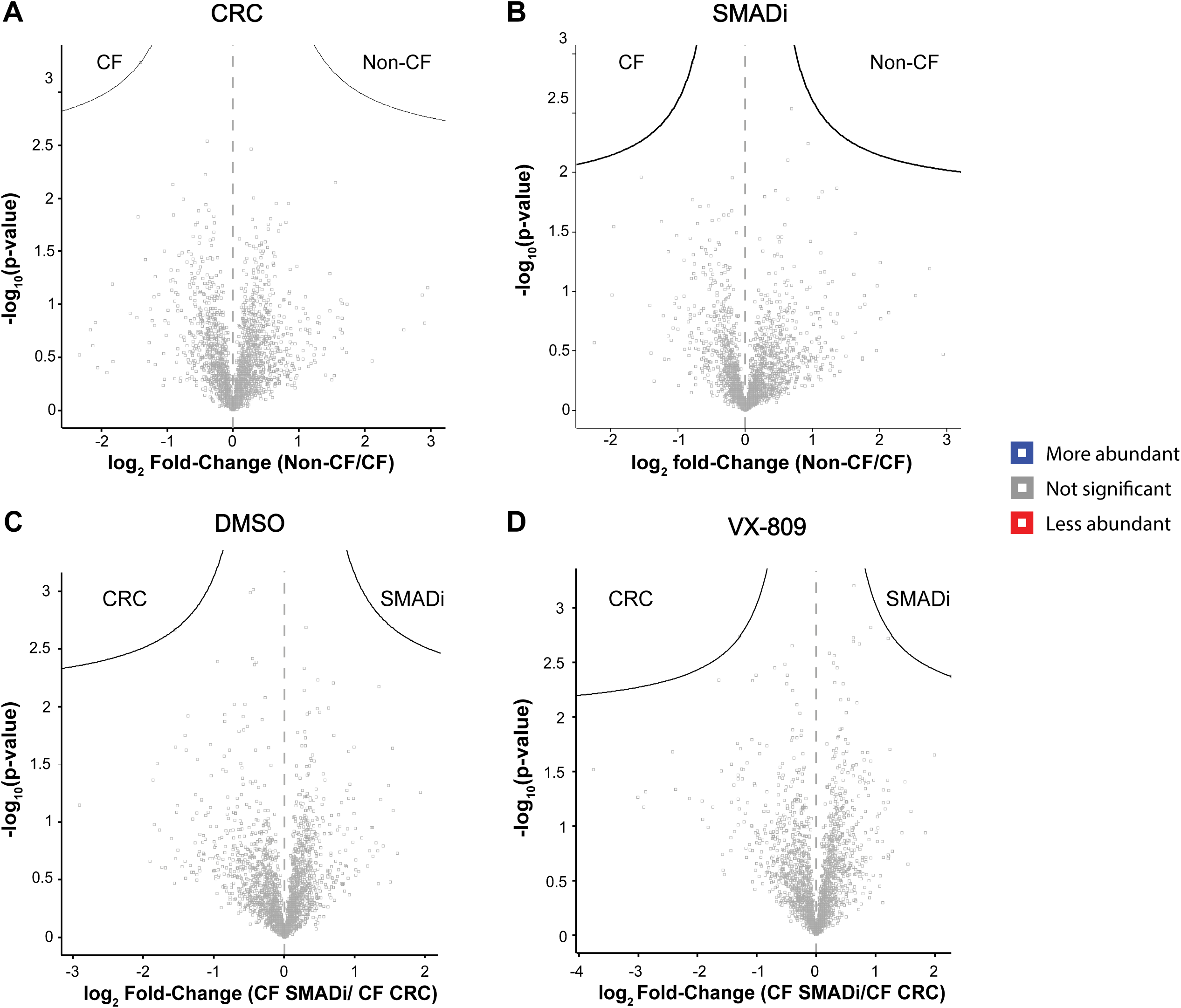
Global proteomic profiles of seven matched F508del/F508del CF and two non-CF CRC^ALI^ vs. SMADi^ALI^. Volcano plot showing differential protein abundance between **(A)** Non-CF and CF CRC^ALI^ cultures, **(B)** Non-CF and CF SMADi^ALI^ cultures, CF SMADi^ALI^ and CF CRC^ALI^ cultures with **(C)** no treatment, and **(D)** with VX-809 treatment. Cut-off curves indicate significant hits (q-value < 0.05 and log2-fold change > 1).

### 3.3 Cilia abundance is the same but cilia beating frequency is higher in CRC^ALI^ than SMAD^ALI^

Both ALI models exhibit functional micro-physiological processes, including beating cilia and the ability to secrete mucus. Positive immunoreactivity of acetylated tubulin (ciliated cell marker) (green; **Fig 4A, Fig S1A**) and MUC5AC (secretory goblet cell marker) were detected at the apical cell surface for both cultures. No skewing towards a specific secretory or more ciliated phenotype was apparent. We confirmed motility of cilia by cilia beat frequency (CBF) measurements. We observed inter-donor heterogeneity in CBF measurements, ranging between 3.72 to 10.05 Hz (**Fig 4B**). CBF values of CF-CRC^ALI^ were significantly higher compared to CF SMADi^ALI^ (7.58 ± 0.24 vs. 6.29 ± 0.32 Hz; P ≤ 0.05) under basal conditions and when cultures were treated with VX-809 (7.64 ± 0.31 vs. 6.22 ± 0.18 Hz; P ≤ 0.001). A similar trend was observed when comparing CRC cultures from the five non-CF donors to their matched SMADi ALI cultures, although statistical significance was not achieved in this group (6.84 ± 0.17 vs. 6.63 ± 0.30 to Hz).

**Fig 4.**
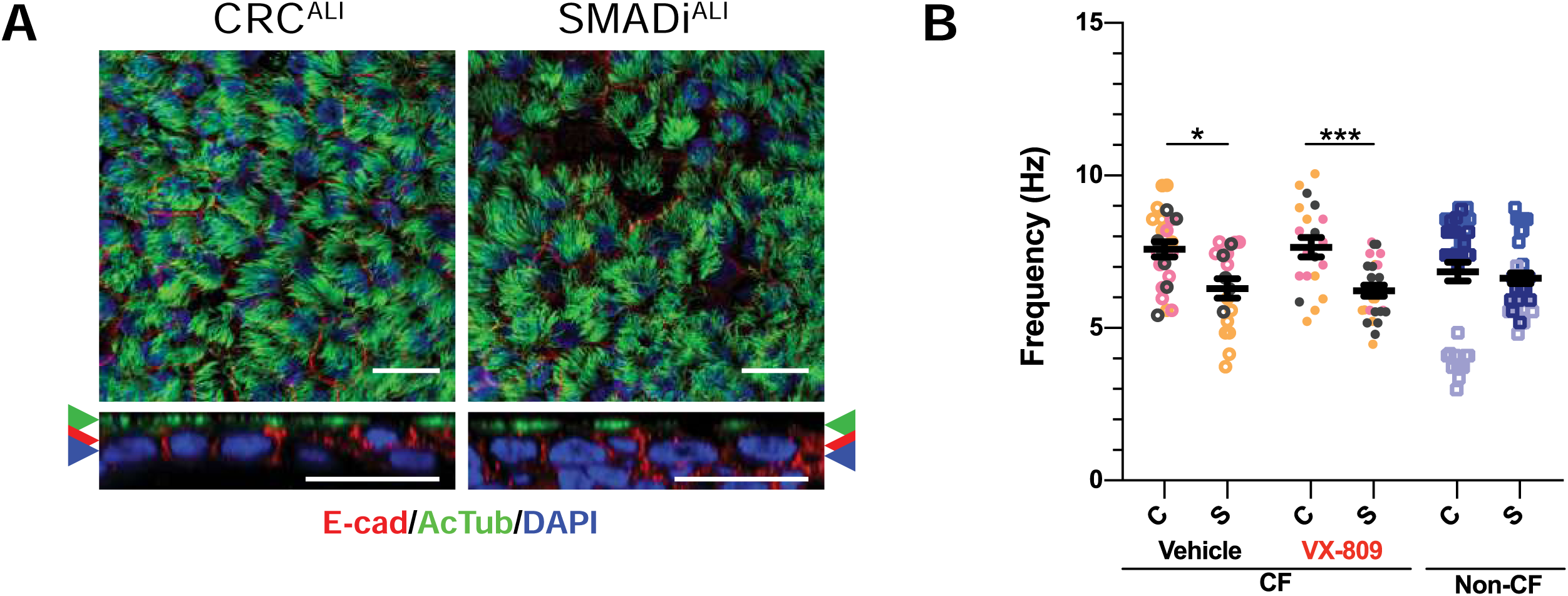
Ciliated cell marker and Cilia beat frequency (CBF) measurement in matched CRC vs. SMADi HNE ALI cultures. **(A)** Ciliated cells acetylated tubulin (green). **(B)** CBF measurements (Hz) in F508del/F508del CFTR (n=3) with and without VX-809 treatment and WT CFTR (n=3). Each coloured dot represents CBF at a field of view. 5-7 different fields of view were sampled per transwell. Dots of the same colour represent the same donor. Open circle represent vehicle (DMSO) treatment and filled circles represent VX-809 treatment for CF donors, and squares indicate non-CF donors. C = CRC, S = SMADi. Error bars represent standard error of the mean (Mean ± SEM). A two-tailed Student’s t-test was used to determine statistical significance. * = P ≤ 0.05 and *** = P ≤ 0.001.

### 3.4 Ion transport functional assessment in matched CRC- and SMADi -ALI cultures

Electrophysiological profiles of CRC^ALI^ and SMADi^ALI^ epithelial cells were created by assessing the function of Amiloride sensitive Epithelial Sodium Channels (ENaC), calcium-activated chloride channel (CaCC) and CFTR mediated Chloride channel (CFTR) from both CF and non-CF donors (**Fig 5, Fig S2, Table S5**). As expected, donor - to - donor variability was evident for all ion channel functions (**Fig S3**).

**Fig 5.**
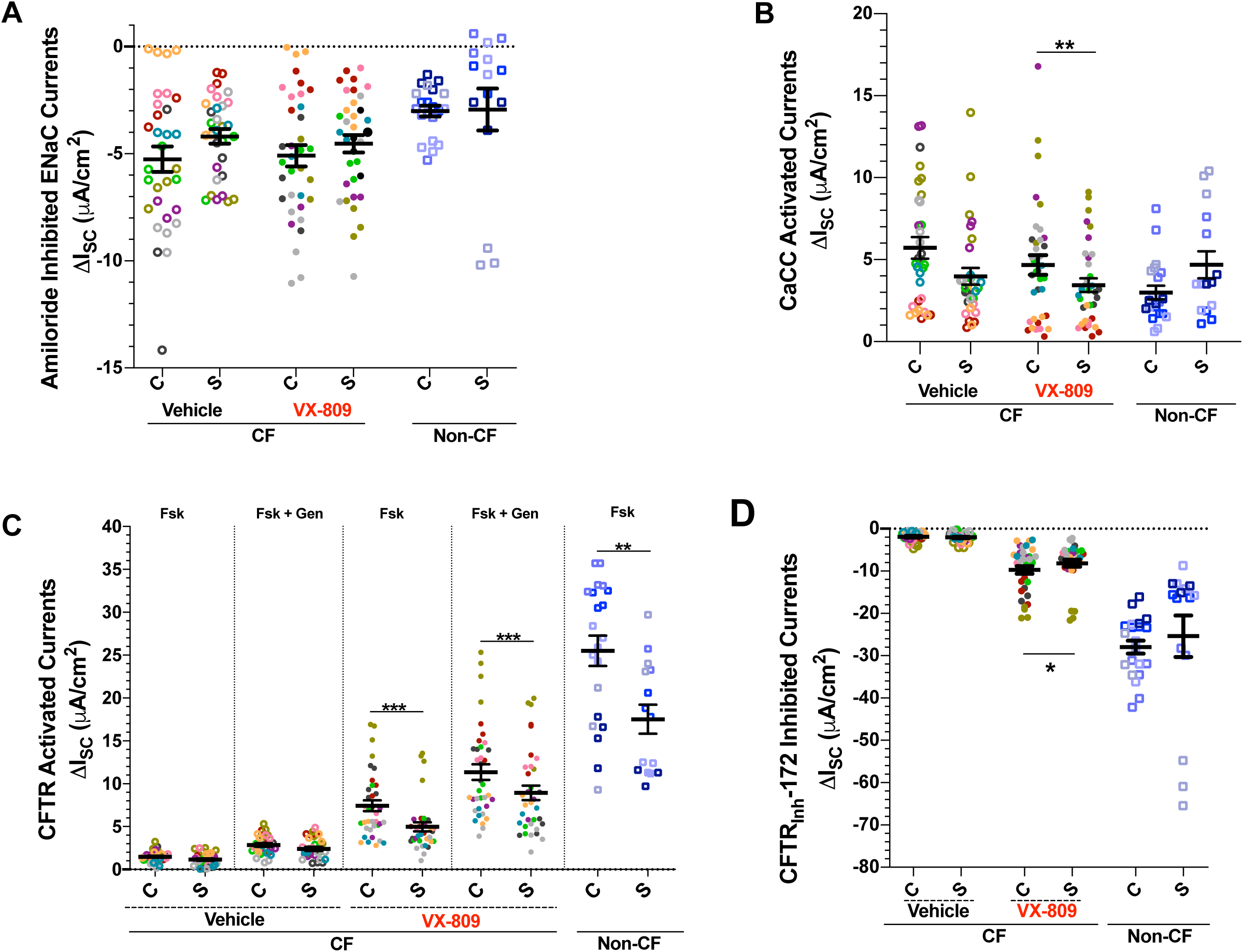
Ion transport measurements in donor matched CRC and SMADi HNE cells grown at Air-Liquid Interface (ALI). Dot plots of mean values of short circuit currents (ΔIsc) **(A)** Amiloride-sensitive ENaC currents, **(B)** ATP-activated CaCC, **(C)** CFTR-Forskolin (Fsk) and Fsk+Genestine (Gen) stimulated and **(D)** inhibited CFTR-dependent activity of F508del/F508del CF (n=9) and WT (n=5) HNE cells. To asses CFTR correction, ALI cell models were pre-incubated with 3 µM VX-809 or 0.03% DMSO (vehicle) for 48 h. Sequential addition of 10 µM Forskolin and 50 µM Genistein in asymmetrical chloride concentration. Each coloured dot represents an individual transwell. Dots of the same colour represent the same donor. Open circle represent vehicle (DMSO) treatment and filled circles represent VX-809 treatment for CF donors, and squares indicate non-CF donors. C = CRC, S = SMADi. Error bars represent standard error of the mean (Mean ± SEM). A two-tailed Student’s t-test was used to determine statistical significance. * = P ≤ 0.05, ** = P ≤0.001 and *** = P ≤ 0.001.

#### 3.4.1 Amiloride-inhibited ENaC currents are similar in CF CRC vs. SMADi HNE ALI cultures

No difference in basal ENaC activity was observed between CRC^ALI^ and SMADi^ALI^ cultures in all donors (**Fig 5A, Fig S3B**). ENaC inhibited currents remained largely similar when treated with VX-809, reinforcing that VX-809 treatment does not modify ENaC inhibited currents.

#### 3.4.2 ATP-stimulated calcium-activated Cl^-^ (CaCC) currents in CRC vs. SMADi HNE ALI cultures

The addition of adenosine triphosphate (ATP) generated transient calcium-activated chloride channel (CaCC) currents (ΔI_sc-ATP_). Both CRC and SMADi cultures displayed variability in ΔI_sc-ATP_ values while the mean average across all cultures were largely comparable, between 3.00 to 5.72 µA/cm^2^ (**Fig 5B, Fig S3C**). VX-809-treated CF SMADi^ALI^ cultures exhibited significantly lower ATP-activated currents compared to the paired CRC ^ALI^ cultures (**Fig 5B**). The difference was significant, but modest (P ≤ 0.05). No disparity was observed in the corresponding non-treated CF CRC ^ALI^ vs. SMADi ^ALI^ cultures, as well as both cultures in non-CF cultures.

#### 3.4.3 Baseline CFTR-mediated Cl^-^ currents in CRC is higher than SMADi HNE ALI cultures

Apical CFTR localisation was confirmed in matched CRC^ALI^ and SMADi^ALI^ cultures derived from a non-CF donor (**Fig S4-Donor 11**). To assess baseline CFTR mediated Cl^-^ secretion, cAMP agonist forskolin was used to activate CFTR Cl^-^ channel, followed by potentiation with genistein. In CF CRC^ALI^ and SMADi^ALI^ cultures, forskolin-induced currents (ΔI_sc-Fsk_) were negligible (**Fig 5C, Fig S3D**). This observation is consistent with previous F508del-CFTR reports of minimal residual function (11, 21). Potentiation of CF ALI cultures with genistein increased forskolin-stimulated currents by nearly 2-fold, up to 2.86 ± 0.20 and 2.40 ± 0.29 µA/cm^2^, respectively (**Fig 5C**).

As anticipated, irrespective of the expansion method, CF ALI cultures had significantly lower CFTR basal activity to the non-CF cultures (**Figure 5C, Fig S2**). In non-CF cultures, forskolin-induced CFTR-mediated currents (ΔI_sc-Fsk_) were significantly higher in the CRC^ALI^ (25.50 ± 1.77 µA/cm^2^) compared to their matched SMADi^ALI^ cultures (17.53 ± 1.69 µA/cm^2^) (P ≤ 0.01; **Fig 5C**).

#### 3.4.4 VX-809-rescued CFTR-mediated Cl^-^ currents in SMADi CF HNE ALI cultures is significantly lower compared to CRC

Treatment of CF CRC^ALI^ and SMADi^ALI^ cultures with VX-809 led to significant correction of CFTR function from all CF donors (**Fig 5C, Fig S2A**). Forskolin-induced currents (ΔI_sc-Fsk_) in CRC^ALI^ increased by 5-fold (from 1.48 ± 0.12 to 7.43 ± 0.63 µA/cm^2^) with VX-809 treatment, and these were further enhanced by genistein by 1.5-fold (up to 11.35 ± 0.90 µA/cm^2^). In contrast, CFTR rescue in the SMADi^ALI^ cultures was significantly smaller with or without genistein potentiation (**Fig 5C**). ΔI_sc-Fsk_ values in SMADi ALI cultures increased by 4-fold (1.16 ± 0.14 to 4.97 ± 0.52 µA/cm^2^) following VX-809-treatment, which were further potentiated 1.8-fold (8.94 ± 0.85 µA/cm^2^) with genistein.

To confirm currents were mediated by the CFTR chloride channel, we used CFTR-specific inhibitor (CFTR_Inh_-172), which almost completely abolished the currents evoked by forskolin + genistein in both cultures, with trends that mirror those of total stimulated CFTR-dependent currents (**Fig 5D**).

## 4. DISCUSSION

Airway epithelial cell models are becoming important pre-clinical tools for personalised CF medicine. Therefore, a better understanding of the impact of culture expansion techniques on CFTR function is warranted. We focused on F508del-CFTR as this is the most prevalent mutation in the CFTR gene. 90% of patients with CF have at least one F508del mutation (22). Two expansion methods that were shown to yield more cells than the methodology using conventional Bronchial Epithelial Cell Growth Medium (BEGM) were tested. At the expansion phase, the morphology of CRC cells were distinct from the SMADi cells, although both had the typical epithelial cobblestone phenotype. In general, the SMADi monolayer presented a more neatly organised cobblestone morphology than the CRC counterpart. These appearances were consistent with those reported in previous studies (16, 19). Our studies showed that the population doubling was significantly higher in CRC HNE compared to SMADi HNE, although, the rate of growth of both cultures were largely similar.

When CRC^ALI^ and SMADi^ALI^ were differentiated using the same protocol, no significant difference in markers of mucociliary differentiation was observed. Despite similar cilia abundance between the two cultures, a significantly lower CBF was observed in the SMADi^ALI^ when compared to the CRC^ALI^. TGF-β has been shown to reduce CBF and ASL volume in CF bronchial epithelial cells, although the role of CFTR was not interrogated in the same studies (23). TGF-β/SMAD/BMP signalling pathways directly regulate CFTR function and biogenesis, as well as epithelial cell behaviour (proliferation, differentiation) (24-27). Potentially, TGF-β/SMAD/BMP signalling has a regulatory role in the mucociliary clearance in a similar manner to the regulation of the CFTR function, although the exact mechanism requires further investigation.

Despite the lack of observed differences in the tight junction (ZO-1) and adherence junction (e-cad) distribution, SMADi^ALI^ cultures displayed ∼1.2-fold higher transepithelial electrical resistance (TEER) values compared to CRC cultures. This indicates neither expansion culture method compromises epithelial tight junction formation after differentiation. This is important for CF research as electrically tight epithelia is pre-requisite to allow for CFTR function measurement (Ussing chamber) (20, 28).

VX-809 is the corrector agent in the first drug (Orkambi) approved for clinical treatment of individuals with homozygous F508del CFTR genotype. It acts by correcting CFTR protein folding and trafficking. A routine *in vitro* assay to characterise CFTR activity in F508del uses treatment with corrector VX-809 to test the rescue of CFTR. 48h treatment with VX-809 demonstrated detectable changes in CFTR function across both the expansion culture types. Donor-to-donor heterogeneity in VX-809 rescue was also seen. CF SMADi cultures evidenced a marked decrease in CFTR mediated Cl^-^ currents compared to CRC^ALI^ cultures. Furthermore, our non-CF SMADi^ALI^ cultures exhibited severe down-regulation of CFTR activity compared to CRC ^ALI^ cultures. Because both cultures were subjected to identical differentiation conditions, any disparity observed in the differentiated CRC^ALI^ and SMADi^ALI^ cultures would be attributed to the distinct expansion conditions. We hypothesise that the attenuated CFTR activity in SMADi^ALI^ cultures could be as a result of a epigenetic remodelling, perhaps a carry-over effect of one or a combination of SMAD inhibitors, used during the expansion phase. Epithelial ion channel function has been shown to be sensitive to epigenetic changes, even when gross differentiation potential is preserved (18, 29).

The (ir)reversibility of the SMAD inhibitors effects on CFTR have not been reported. A83-01 acts through blocking Smad2/3 activity while DMH1 blocks Smad 1/5/8 (30, 31). This combination abrogates the actions of all SMAD family proteins, the transcription factors downstream of the growth factor and cytokine TGF-β (32). The relationship of TGF-β and CFTR is complex. TGF-β is a well-known genetic modifier of CF lung disease progression with certain polymorphisms of the TGF-β having been associated with a more severe CF phenotype (33, 34). TGF-β expression in human lung tissue does not appear de-regulated in CF when compared to healthy control, but TGF-β signalling (phosphorylated Smad2 expression) was found to be upregulated (35). In CF patients, the levels of TGF-β in their blood serum increased during disease exacerbation and *P. aeruginosa* infection (36). In addition, TGF-β significantly down-regulates the level of CFTR mRNA and cAMP-dependent CFTR currents in cultures established from non-CF nasal polyps (37). TGF-β also inhibited VX-809 corrected F508del CFTR in colonic and human airway epithelial cells (24, 38, 39). Given the evidence for the negative regulatory role of TGF-β on CFTR, we expected that TGF-β inhibitors A83-01 and DMH1 effect would manifest as increased CFTR activity in the SMADi^ALI^ cultures.

With TGF-β pleiotropic activity (32), it is possible for inhibition of TGF-β/SMAD pathways to act upon and down-regulate the CFTR function. This is entirely plausible given Smad3 expression is reduced in nasal epithelial tissues of 18 CF patients compare to five healthy controls despite enhanced TGF-β signalling reported in separate studies (35, 40). The reduced Smad3 levels in CF airways can explain the relative insensitivity/smaller disparity in CFTR function elicited in the CF CRC vs. SMADi cultures when compared to non-CF cultures. On the other hand, we did not observe any significant changes in the basal activity of ATP activated CaCC currents between the two cultures from the CF and non-CF individuals. We thus suggest that SMAD inhibitory effect is likely to be CFTR-specific.

It appears that the alteration of cellular physiological function *in vitro* occurs more easily than structural and differentiation potential. We cannot ascertain which one of the two culture expansion methods studied here correlates more with the *in vivo* CFTR function, since none of the donors in this study were receiving CFTR modulator therapy. It is necessary to recognise the limitations of *in vitro* cultures as pre-clinical models for CF. Culture conditions significantly influence CFTR activity. This could lead to false conclusions when data from various labs are compared against each other. With the current variety of techniques in use, it is important to compare like for like and report a patient’s cell model’s response to drugs against technique/lab specific references. Moving forward, a standardised protocol to expand and differentiate patient airway epithelial cells is needed across CF labs to ensure consistency in *in vitro* data and, ultimately, translation of clinical care to patients.

## Abbreviations

CF: (Cystic Fibrosis),
CFTR: (Cystic Fibrosis Transmembrane Conductance Regulator),
ALI: (air liquid interface),
ATP: (Adenosine tri phosphate),
HBE: (human bronchial epithelial cell),
HNE: (human nasal epithelial cell),
CRC: (conditionally reprogrammed cultures),
ROCK: (Rho-associated protein kinase),
P: (passage),
TEER: (Trans-epithelial electrical resistance),
TGF-β: (Transforming growth factor-beta),
I_sc_: (Short circuit current),
CaCC: (Calcium activated chloride channels),
CBF: (Ciliary beating frequency)

## Acknowledgements

We thank the study participants and their families for their contributions. We appreciate the assistance from Sydney Children’s Hospitals (SCH) Randwick respiratory department in organization and collection of patient biospecimens – special thanks to Dr John Widger, Dr Yvonne Belessis, Leanne Plush, Amanda Thompson and Rhonda Bell. This work was supported by Australian National Health and Medical Research Council (NHMRC_APP1188987), Cystic Fibrosis Australia, Sydney Children Hospital Foundation and Luminesce Alliance Research grants. We thank Dr. John R. Riordan (University of North Carolina – Chapel Hill) and Cystic Fibrosis Foundation for providing anti-CFTR antibodies #570, 596, 769, 450 and 528.

## Author contributions

Funding acquisition: SAW, AJ. Conceptualization, design and supervision: SAW. Consent and biospecimens collection: LKF, AJ. Patient samples processing: NT, NTA and SLW. Experimental work, data analysis and interpretation: NTA, SLW, EP, IS, LZ, AC and SAW. Manuscript writing: SLW, NTA and SAW with intellectual input from all authors.

## Supplementary Files

### SUPPLEMENTARY MATERIALS and METHODS

#### Study Participants

Paediatric CF patients with homozygous F508del-CFTR (n=9) and non-CF controls (n=5) were included in this study (**Table S1**). No study participant was on CFTR modulator therapy prior to sample acquisition. This study was approved by the Sydney Children’s Hospital Ethics Review Board (HREC/16/SCHN/120), and written informed consent was obtained from the guardians of all participants.

#### Nasal airway cell procurement and processing

Primary human nasal epithelial (HNE) cells were obtained through cytology brushing of the nasal inferior turbinate of CF participants during annual surveillance clinic or bronchoscopy. Nasal brushings for non-CF participants were collected during elective bronchoscopy or non-respiratory related investigative procedures. HNE cells were dislodged from cytology brushes by gentle vortexing in collection media. Cells were pelleted at 300 g for 5 min at 4°C, resuspended with the expansion culture media specified below, and passed through a 40 µm cell strainer (Sigma CLS431750) to generate single-cell suspension. Cells were then seeded equally into flasks for serial expansion using two culture methods, conditionally reprogrammed cell (CRC) culture and feeder-free dual SMAD inhibition (SMADi) culture.

#### Conditionally reprogrammed cell (CRC) expansion culture – co-culture method

Isolated HNE cells were co-cultured with irradiated NIH/3T3 feeder cells (culture details below) in F-media containing Rho kinase (ROCK) inhibitor, Y-27632 as described previously (14). HNE cells were added to collagen I-coated flask pre-seeded with mitotically arrested NIH/3T3 feeder cells in F-media (**Table S2**), and media change was performed every second day until 80-90% confluence. Cells were dissociated using a differential trypsin method. Cells were neutralized with trypsin neutralizing solution (Lonza CC-5034), and cell count was performed using Countess II automated cell counter (Thermo Fisher Scientific, Waltham, MA) according to the manufacturer’s instructions and then seeded for ALI cultures or cryopreserved.

#### NIH/3T3 feeder cell culture and irradiation

NIH/3T3 mouse embryonic fibroblast cell line was cultured at 37°C, 5% CO_2_ in DMEM (Life Technologies 11965-092) supplemented with 10% FBS, 100 U/ml penicillin and 100 µg/ml streptomycin. When indicated for gamma-irradiation, cells at 70-80% confluency were trypsinized, and pelleted cells were resuspended in fresh culture media. The cell suspension was exposed to 30 Gy gamma-irradiation (J.L. Shepherd & Associates, San Fernando, CA) and then seeded into flasks coated with collagen I (PureCol; Advanced Biomatrix 5005) at a density of 5000 cells/cm^2^.

#### Dual SMAD inhibition (SMADi) expansion culture – serum and feeder-free method

Isolated HNE cells were seeded into a collagen I-coated flask, as described previously with minor modifications (16). Cells were cultured in Bronchial Epithelial Cell Medium (BEpiCM) (ScienCell Research Laboratories 3211) supplemented with 1 µM A83-01 (Tocris Bioscience 2939), 1 µM DMH1 (Selleckchem S7146), 3.3 nM EC23 (Enzo Life Sciences BML-EC23-0500) and 10 µM Y-27632. Media change was performed every second day until 80-90% confluence. Cells were dissociated with trypsin/EDTA (Lonza CC-5034) for 5-7 min at 37°C and neutralized with trypsin neutralizing solution (Lonza CC-5034). Cell count was performed using Countess II automated cell counter according to the manufacturer’s instructions and then seeded for ALI cultures or cryopreserved.

#### Mucociliary differentiation at air-liquid interface (ALI)

Passage one, donor-matched HNE CRC- and SMADi-expanded cells were seeded on Transwell 6.5mm, 0.4µm pore polyester membrane inserts (0.33cm^2^ area; Sigma CLS3470) pre-coated with collagen I (PureCol; Advanced Biomatrix 5005), at a density of 150,000 cells/insert till confluence. Once confluent under submerged conditions (approximately 4-5 days), cells were switched to air-liquid interface (ALI) culture condition, whereby apical media was removed, and PneumaCult ALI differentiation media (STEMCELL Technologies, 05001) was added to the basolateral compartment only. Basal media change was performed every second day for 3-4 weeks. Beating cilia (ciliogenesis) and mucus production were monitored using light microscopy. The apical surface was washed with warm phosphate buffered saline (PBS) once a week to remove excess mucus. After 21-25 days, Ussing chamber measurements were carried out with the resistance values above 200 Ω.cm^2^. ALI cultures established from HNE CRC- and SMADi-expanded cells were incubated (basolateral side) with 3 µM VX-809 (Selleckchem S1565), a CFTR corrector compound or DMSO 0.1%v/v (vehicle) for 48 h.

#### Short circuit current measurements in Ussing chambers

HNE ALI Transwell inserts were mounted in circulating Ussing chambers (VCC MC8 multichannel voltage/current clamp; Physiologic Instruments). Three to six transwells were tested per donor per condition. The short circuit current I_SC_ and transepithelial resistance were measured under voltage-clamp conditions. For I_SC_ recordings, a basolateral-to-apical Cl^-^ secretory gradient was created using asymmetric chloride (Cl^-^) Ringer’s buffer. The basolateral Cl^-^ solution contained (mM): 145 NaCl, 3.3 K_2_HPO_4_, 1.2 CaCl_2_, 1.2 MgCl_2_, 10 HEPES and 10 glucose, pH 7.4. The composition of apical low-Cl^-^ solution was the same apart from the replacement of NaCl with equimolar Na-gluconate. Ringer’s buffers were continuously gassed with 95% O_2_-5% CO_2_ and maintained at 37°C. Following a 30min stabilisation period cells were treated with pharmacological compounds (in order): 100 µM amiloride (apical) to inhibit epithelial sodium channel (ENaC)-mediated Na^+^ flux, 10 µM forskolin (basal) to induce cAMP activation of CFTR, 50 µM genistein (apical) to further potentiate cAMP-activated currents, 30 µM CFTR_inh_-172 (apical) to inhibit CFTR-specific currents and 100 µM ATP (apical) to activate purinergic calcium-activated chloride currents. Cumulative changes of I_sc_ in response to forskolin and genistein (ΔI_sc-Fsk+Gen_) was used as an indicator of maximum CFTR function. Data recordings were acquired using Acquire and Analyze (version 2.3) software (Physiologic Instruments).

#### Sample preparation for Mass Spectrometry

ALI differentiated HNEC cultures untreated or treated with VX-809 were harvested for mass spectrometry. Total protein was extracted by homogenizing the cells in RIPA buffer (Life Technologies 89900) containing protease inhibitor cocktail (Sigma 11836153001). Samples were sonicated using the Bioruptor Pico (Diagenode B01060010) for a total of 10 min using a 30 sec on/off cycle at 4°C. Protein concentrations were determined using the 2-D Quant kit (GE Life Sciences 80648356). Samples were reduced (5 mM DTT, 37C, 30 min), alkylated (10 mM IA, RT, 30 min) then incubated with trypsin at a protease:protein ratio of 1:20 (w/w) at 37°C for 18 h, before being subjected to SCX clean-ups (Thermo Fisher, SP341) following manufacturer’s instructions. Eluted peptides from each clean-up were evaporated to dryness in a SpeedVac and reconstituted in 20 µL 0.1% (v/v) formic acid, 0.05% HFBA and 2% acetonitrile.

#### Mass Spectrometry

Proteolytic peptide samples were separated by nanoLC using an Ultimate nanoRSLC UPLC and autosampler system (Dionex, Amsterdam, Netherlands. A micro C18 precolumn with H_2_O:CH_3_CN (98:2, 0.1 % TFA) at 15 µL/min and a fritless nano column (75 µm x 15 cm) containing C18-AQ media (Dr Maisch, Ammerbuch-Entringen Germany) was used to concentrate and desalt samples. Peptides were eluted through a linear gradient of H_2_O:CH_3_CN (98:2, 0.1 % formic acid) to H_2_O:CH_3_CN (64:36, 0.1 % formic acid) at 200 nL/min over 30 min. Eluted peptides were ionized using positive ion mode nano-ESI by applying 2000 volts to a low volume titanium union with the tip positioned ∼0.5 cm from the heated capillary (T=275°C) of a Tribrid Fusion Lumos mass spectrometer (Thermo Scientific, Bremen, Germany).

A survey scan *m/z* 350-1750 was acquired in the orbitrap (resolution = 120,000 at *m/z* 200, with an accumulation target value of 400,000 ions) and lockmass enabled (*m/z* 445.12003). Data dependant acquisition was used to sequentially select peptide ions (>2.5×10^4^ counts, charge states +2 to +5) for MS/MS, with the total number of dependent scans maximized within 2 sec cycle times. Product ions were generated via higher energy collision dissociation (collision energy = 30; maximum injection time = 250 milliseconds; MS_n_ AGC = 5×10^4^; inject ions for all available parallelizable time enabled) and mass analyzed in the linear ion trap. Dynamic exclusion was enabled and set to: n times =1, exclusion duration 20 seconds, ± 10ppm. Mass spectrometry data are available at the ProteomeXchange Consortium via the PRIDE partner repository with the dataset identifier PXD018386 (https://www.ebi.ac.uk/pride/archive/projects/PXD018386). Full list of identified proteins and differentially abundant protein analysis is available upon request.

#### Protein identification, quantification and statistical analysis

LC-MS/MS raw files were analysed using the MaxQuant software suite (version 1.6.2.10.43) (41). Sequence database searches were performed using Andromeda (42). Label-free protein quantification was performed using the MaxLFQ algorithm (43). Delayed normalizations were performed following sequence database searching of all samples with tolerances set to ±4.5 ppm for precursor ions and ±0.5 Da for peptide fragments. Additional search parameters were: carbamidomethyl (C) as a fixed modification; oxidation (M) and N-terminal protein acetylation as variable modifications; and enzyme specificity was trypsin with up to two missed cleavages. Peaks were searched against the human Swiss-Prot database (August 2018 release), which contained 20333 sequences with the minimum peptide length set as 7. MaxLFQ analyses were performed using default parameters with “fast LFQ” enabled. Protein and peptide false discovery rate (FDR) thresholds were set at 1% and only non-contaminant proteins identified from ≥2 unique peptides were subjected to downstream analysis.

Statistical analyses of protein abundances were performed with Perseus (version 1.6.5.0) platform. Hits only identified by site, reverse hits and potential contaminants were filtered out. Only proteins that were present in 3 out of 6 replicates were retained. Protein intensities were log_2_-transformed. Missing values were added from normal distribution. Student’s t-tests were performed with Benjamini-Hochberg correction to identify differentially abundant proteins (q-value < 0.05). Volcano plots were constructed using t-test with 250 randomizations.

#### Cilia beating frequency measurement

Cilia beating were imaged in transmission light modality using ORCA-Flash 4.0 sCMOS camera (Hamamatsu Photonics, Shizuoka Pref., Japan), connected to Zeiss Axio Observer Z.1 inverted microscope (Carl Zeiss, Jena, Germany). Images were acquired serially at 334 frames per second (fps) with 3 ms exposure time, on a EC Plan-Neofluar 20x/0.5 Ph2 M27 dry objective (512 × 512 pixels field of view; 0.325 µm x 0.325 µm per pixel). Five to seven time-series were sampled at random from triplicate filters per treatment condition. Imaging was performed at 37°C, 5% CO_2_ to mimic the physiological environment. To extract cilia beating spectra and corresponding beating frequency, image series were analysed using a custom-built script in Matlab (MathWorks, Natick, MA). Briefly, imported image series were filtered to remove the immobile component in each pixel by subtracting the temporal average image, or the DC component. This step ensures that all mucus and other immobile structures producing high scattering in the transmitted image series are excluded from analysis, and only the moving (mobile) parts of the images are processed for spectra and beating frequency recovery. The temporal spectrum for each pixel in the image series was then computed using the Fast Fourier Transform (FFT) algorithm. The peaks in the spectrum indicate frequencies at which the temporal pixel intensity oscillates. The average spectrum per field of view was calculated using the average of all the single pixel spectra. The dominant frequency (a frequency with the highest peak) was then identified using the Matlab function ‘findpeaks’.

#### Whole mount immunofluorescence

After completion of short-circuit current measurement, HNE ALI cultures were washed with PBS at room temperature for three times, 5 min each (Sigma D8662) and fixed. Immunostaining was performed only on cells pre-incubated with DMSO (vehicle) for 48 h or with no pre-incubation. Different fixative solutions were used depending on the target protein. For staining of mucociliary differentiation markers (**Table S3**), cells were fixed in 4% paraformaldehyde for 15 min at room temperature or in methanol-acetone (1:1) for 15 min at -20°C, and then permeabilized with 0.5% Triton-X in PBS on ice for 10min. For CFTR staining, cells were fixed in ice-cold acetone for 15min at -20°C with no permeabilization step. Fixed, permeabilized cells on transwell membranes were rinsed with PBS 3 times and blocked using IF buffer (0.1% BSA, 0.2% Triton and 0.05% Tween 20 in PBS) with 10% normal goat serum for 1 hour at room temperature. Membranes were excised from transwell inserts using a sharp scalpel (size 11) at the end of the blocking step and cut into 2 or 3 equal pieces. Cells were then incubated in primary antibodies (**Table S3**) overnight at 4**°**C on SuperFrost Plus slides (Thermo Fisher Scientific, Waltham, MA). Cells were washed with IF buffer (3X, 5 min each) and incubated with Alexa Fluor conjugated secondary antibodies (Table S**3**) for 1 hour at room temperature. Cells were washed with IF buffer (3X, 5min each) and mounted with Vectashield hardset antifade mounting medium containing DAPI (H-1500; Vector Laboratories, Burlingame, CA). Images were acquired using Leica TCS SP8 DLS confocal microscope (Leica Microsystems, Wetzlar, Germany), 63x/1.4 oil immersion objective. Images were then processed using ImageJ software (National Institute of Health, Bethesda, MD)

#### Statistical analysis

Data were presented as means ± standard error of the mean (SEM). The two-tailed student’s t-test was used to determine the differences between the two groups. Statistical analysis was performed with GraphPad Prism 8 software (GraphPad Software, San Diego, CA). P < 0.05 was considered to be statistically significant.

---

**Table S1.**
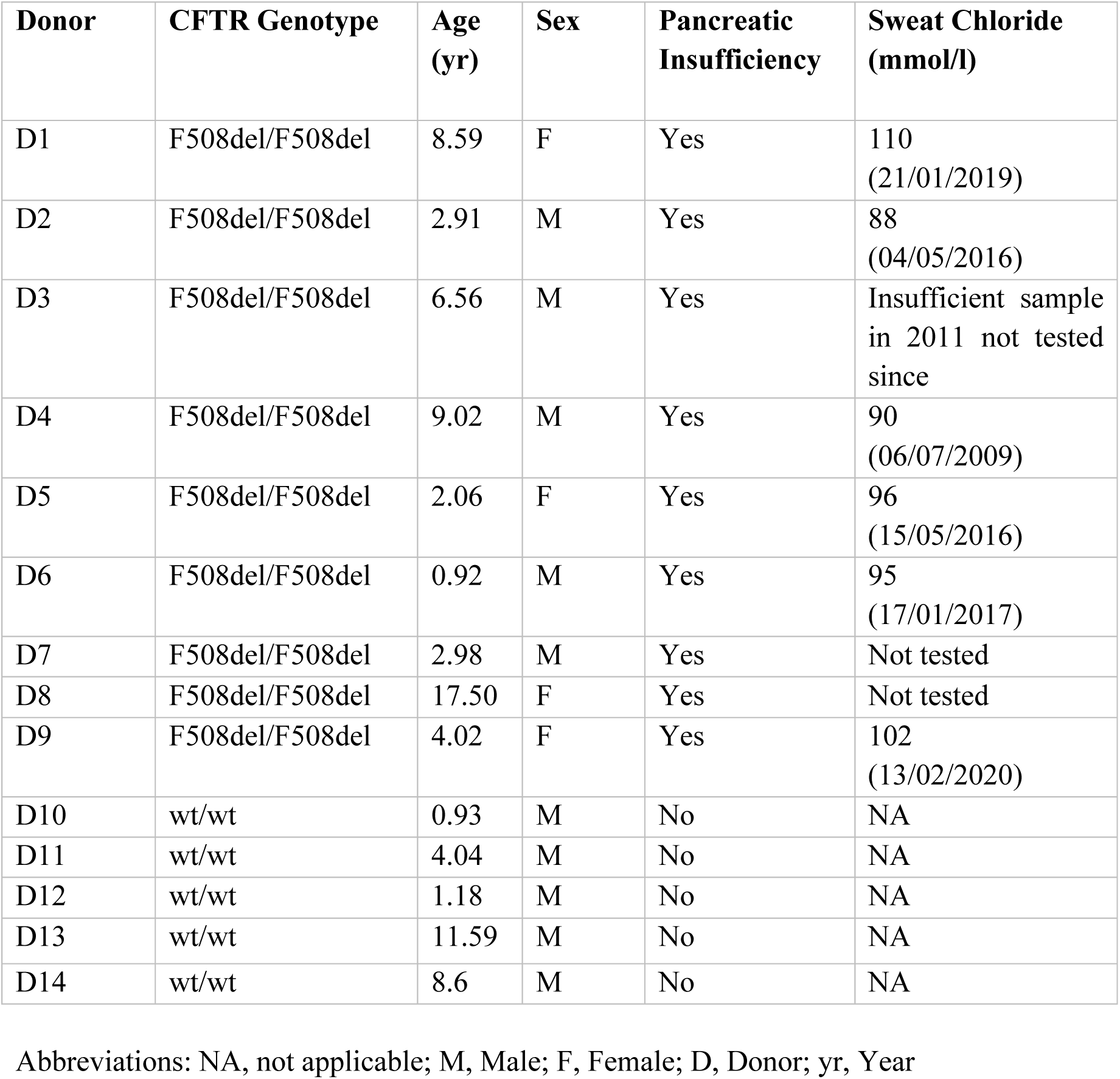
Demographics of study participants from whom HNE cells were cultured.

| Donor | CFTR Genotype | Age (yr) | Sex | Pancreatic Insufficiency | Sweat Chloride (mmol/l) |
| --- | --- | --- | --- | --- | --- |
| D1 | F508del/F508del | 8.59 | F | Yes | 110<br>(21/01/2019) |
| D2 | F508del/F508del | 2.91 | M | Yes | 88<br>(04/05/2016) |
| D3 | F508del/F508del | 6.56 | M | Yes | Insufficient sample<br>in 2011 not tested<br>since |
| D4 | F508del/F508del | 9.02 | M | Yes | 90<br>(06/07/2009) |
| D5 | F508del/F508del | 2.06 | F | Yes | 96<br>(15/05/2016) |
| D6 | F508del/F508del | 0.92 | M | Yes | 95<br>(17/01/2017) |
| D7 | F508del/F508del | 2.98 | M | Yes | Not tested |
| D8 | F508del/F508del | 17.50 | F | Yes | Not tested |
| D9 | F508del/F508del | 4.02 | F | Yes | 102<br>(13/02/2020) |
| D10 | wt/wt | 0.93 | M | No | NA |
| D11 | wt/wt | 4.04 | M | No | NA |
| D12 | wt/wt | 1.18 | M | No | NA |
| D13 | wt/wt | 11.59 | M | No | NA |
| D14 | wt/wt | 8.6 | M | No | NA |

Abbreviations: NA, not applicable; M, Male; F, Female; D, Donor; yr, Year

**Table S2.** Components of CRC expansion media.

| Components | Concentrations | Supplier |
| --- | --- | --- |
| DMEM/Ham's F-12 | 67% | Life Technologies 11330-032 |
| DMEM, high glucose | 33% | Life Technologies 11965-092 |
| Fetal bovine serum (FBS) | 5% | Life Technologies 10082-147 |
| Penicillin-Streptomycin | 1x | Sigma P4333 |
| Hydrocortisone | 0.4 µg/ml | Sigma H0888 |
| Insulin | 5 µg/ml | Sigma I2643 |
| Cholera toxin | 8.4 ng/ml | Sigma C8052 |
| hEGF | 25 ng/ml | Sigma E9644 |
| Adenine | 24 µg/ml | Sigma A2786 |
| Y-27632 | 10 µM | Selleckchem S1049 |

**Table S3.**
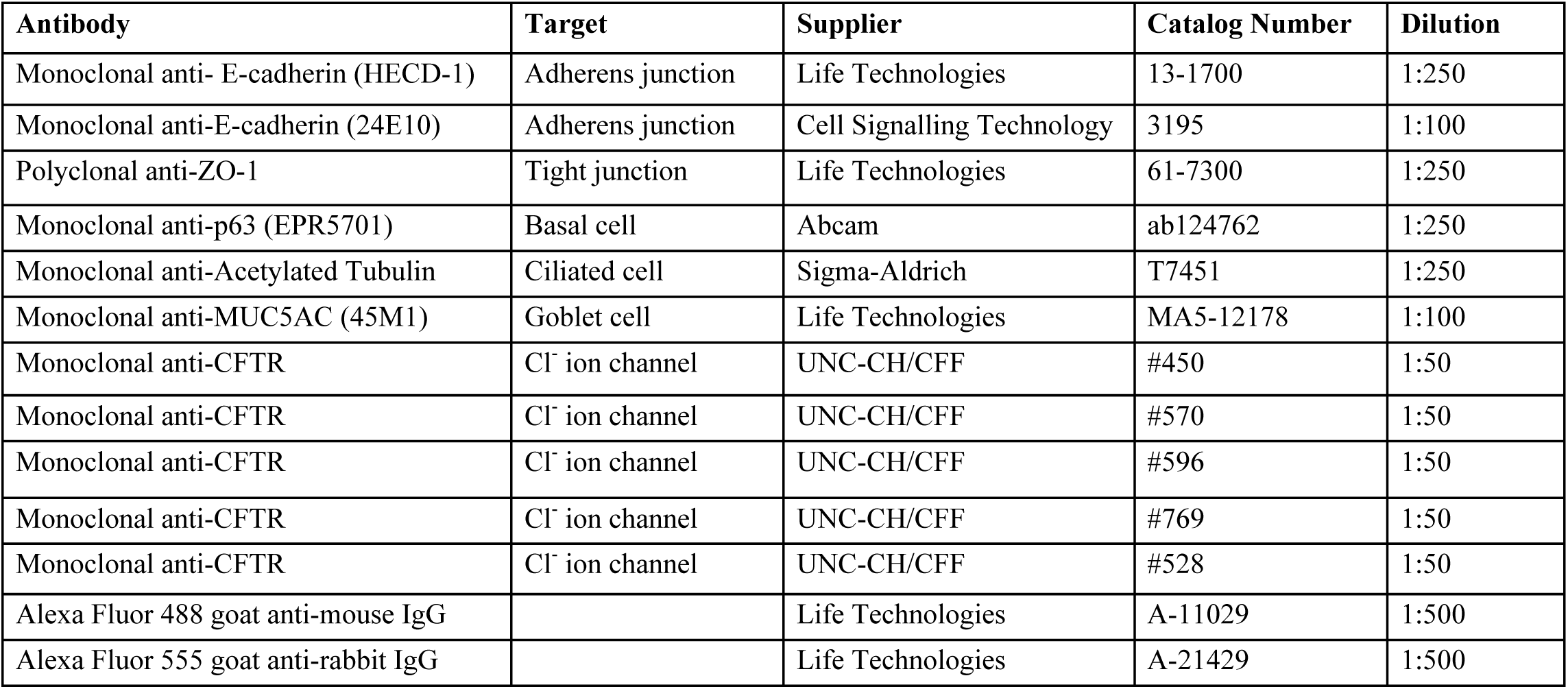
Antibodies used for immunofluorescence characterization.

| Antibody | Target | Supplier | Catalog Number | Dilution |
| --- | --- | --- | --- | --- |
| Monoclonal anti- E-cadherin (HECD-1) | Adherens junction | Life Technologies | 13-1700 | 1:250 |
| Monoclonal anti-E-cadherin (24E10) | Adherens junction | Cell Signalling Technology | 3195 | 1:100 |
| Polyclonal anti-ZO-1 | Tight junction | Life Technologies | 61-7300 | 1:250 |
| Monoclonal anti-p63 (EPR5701) | Basal cell | Abcam | ab124762 | 1:250 |
| Monoclonal anti-Acetylated Tubulin | Ciliated cell | Sigma-Aldrich | T7451 | 1:250 |
| Monoclonal anti-MUC5AC (45M1) | Goblet cell | Life Technologies | MA5-12178 | 1:100 |
| Monoclonal anti-CFTR | Cl<sup>-</sup> ion channel | UNC-CH/CFF | #450 | 1:50 |
| Monoclonal anti-CFTR | Cl<sup>-</sup> ion channel | UNC-CH/CFF | #570 | 1:50 |
| Monoclonal anti-CFTR | Cl<sup>-</sup> ion channel | UNC-CH/CFF | #596 | 1:50 |
| Monoclonal anti-CFTR | Cl<sup>-</sup> ion channel | UNC-CH/CFF | #769 | 1:50 |
| Monoclonal anti-CFTR | Cl<sup>-</sup> ion channel | UNC-CH/CFF | #528 | 1:50 |
| Alexa Fluor 488 goat anti-mouse IgG |  | Life Technologies | A-11029 | 1:500 |
| Alexa Fluor 555 goat anti-rabbit IgG |  | Life Technologies | A-21429 | 1:500 |

**Table S4.** Differential protein abundance analysis of TGF-β/SMAD pathway proteins between both CF and non-CF CRC and SMADi cultures.

| Protein | Log2 fold-change | p-value | q-value |
| --- | --- | --- | --- |
| **CF CRC vs. SMADi** |  |  |  |
| CUL1 | -0.566423098 | 0.00954724 | 0.96373238 |
| MAPK14 | 0.787147458 | 0.10895098 | 0.96373238 |
| SKP1 | -0.354222234 | 0.14252945 | 0.96373238 |
| SMAD2;SMAD3 | -0.296781603 | 0.1856753 | 0.96373238 |
| PPP2R1A | -0.168820635 | 0.28337687 | 0.96373238 |
| PPP2CA | -0.252598 | 0.28830046 | 0.96373238 |
| MAP2K2 | 0.181367811 | 0.31165083 | 0.96373238 |
| ITGB1 | 0.222642326 | 0.33913523 | 0.96373238 |
| MAPK3 | 0.137226931 | 0.51044733 | 0.96373238 |
| RBX1 | 0.224527359 | 0.53252339 | 0.96373238 |
| STRAP | 0.075380961 | 0.55691665 | 0.96373238 |
| HDAC1 | -0.094338799 | 0.61290863 | 0.96373238 |
| MAPK1 | 0.050602595 | 0.74926934 | 0.97586445 |
| SIN3A | -0.070233536 | 0.87339451 | 0.97793369 |
| RHOA;RHOC;ARHA | 0.016871834 | 0.95189139 | 0.98554262 |
| **Non-CF CRC vs. SMADi** |  |  |  |
| PPP2R1A | -0.124732971 | 0.02124631 | 0.99951841 |
| PPP2CA | -0.231428146 | 0.04931977 | 0.99951841 |
| CUL1 | -0.652365685 | 0.21696662 | 0.99951841 |
| RHOA | 0.302098274 | 0.21779762 | 0.99951841 |
| MAPK1 | -0.119151115 | 0.32568562 | 0.99951841 |
| MAPK3 | 0.130068779 | 0.55281491 | 0.99951841 |
| RBX1 | -0.395497322 | 0.6652518 | 0.99951841 |
| SMAD2 | -0.048676491 | 0.87870185 | 0.99951841 |
| SMAD3 | -0.048676491 | 0.87870185 | 0.99951841 |
| SKP1 | 0.054797173 | 0.91473522 | 0.99951841 |
| PPP2CB | 0 | NaN | NaN |
| ROCK1 | 0 | NaN | NaN |
| SMAD4 | 0 | NaN | NaN |
| SP1 | 0 | NaN | NaN |
| CDKN2B | 0 | NaN | NaN |

**Table S5.**
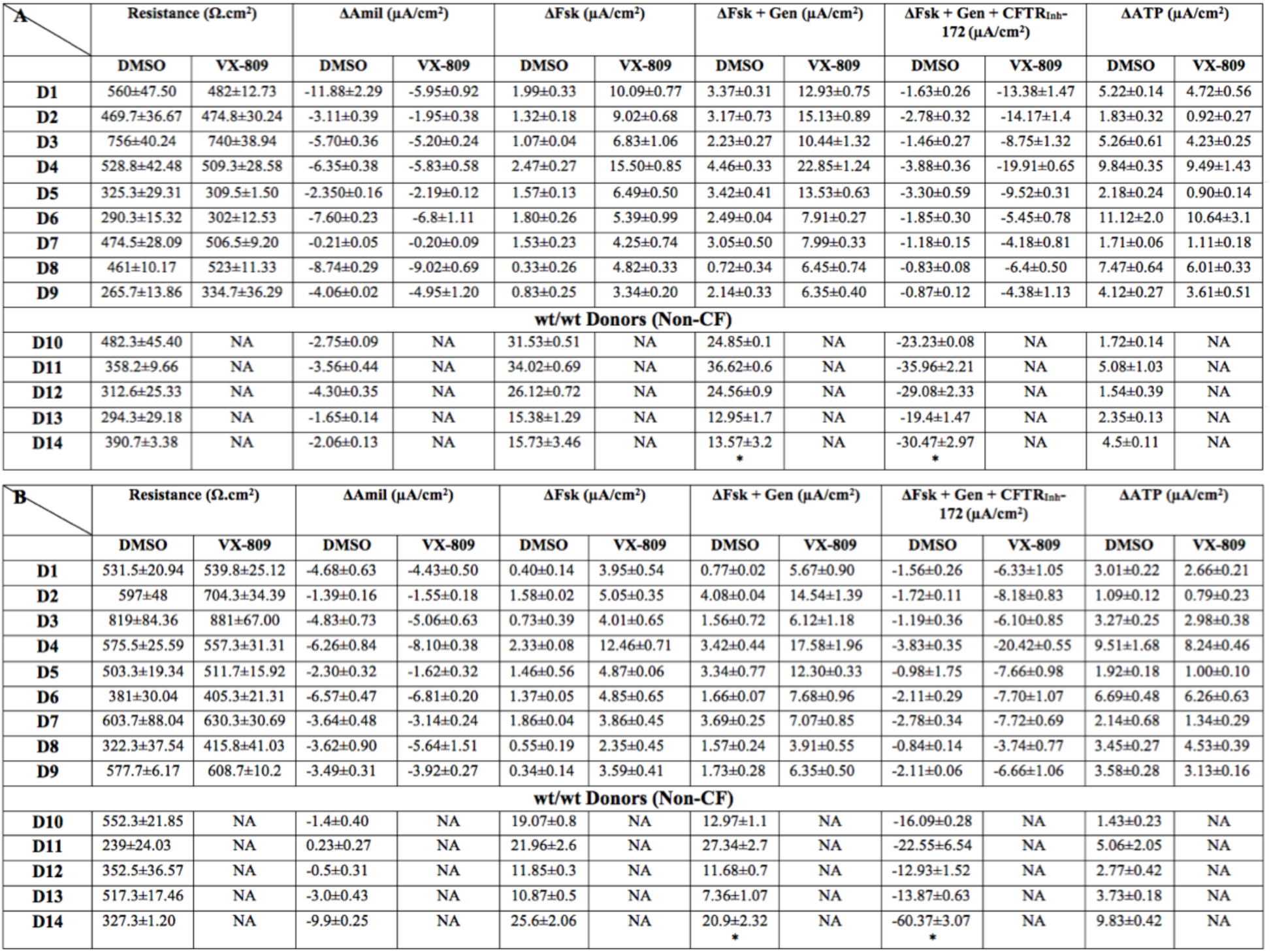
Data for the short-circuit currents and electrophysiological parameters in (A) and (B) SMADi ALI cultures from CF and Non-CF donors. Data represent short circuit current values for Resistance of the monolayers, Amiloride inhibited ENaC currents (ΔAmil), Forskolin stimulated cAMP currents alone (ΔFsk), Genistein potentiatrd currents (ΔFsk + Gen), *VX-770 potentiated currents (ΔFsk+ VX-770), CFTR_Inh_-172 inhibited currents (ΔFsk + Gen + CFTR_Inh_-172), *(ΔFsk + VX-770 + CFTR_Inh_-172), and ATP-achieved currents (ΔATP) (Under DMSO and VX-809 treatment). Values represented are after vehicle or VX-809 (3 μM/48h) treatments; (mean ±SEM); NA = not applicable and D= Donor.

**Fig S1.**
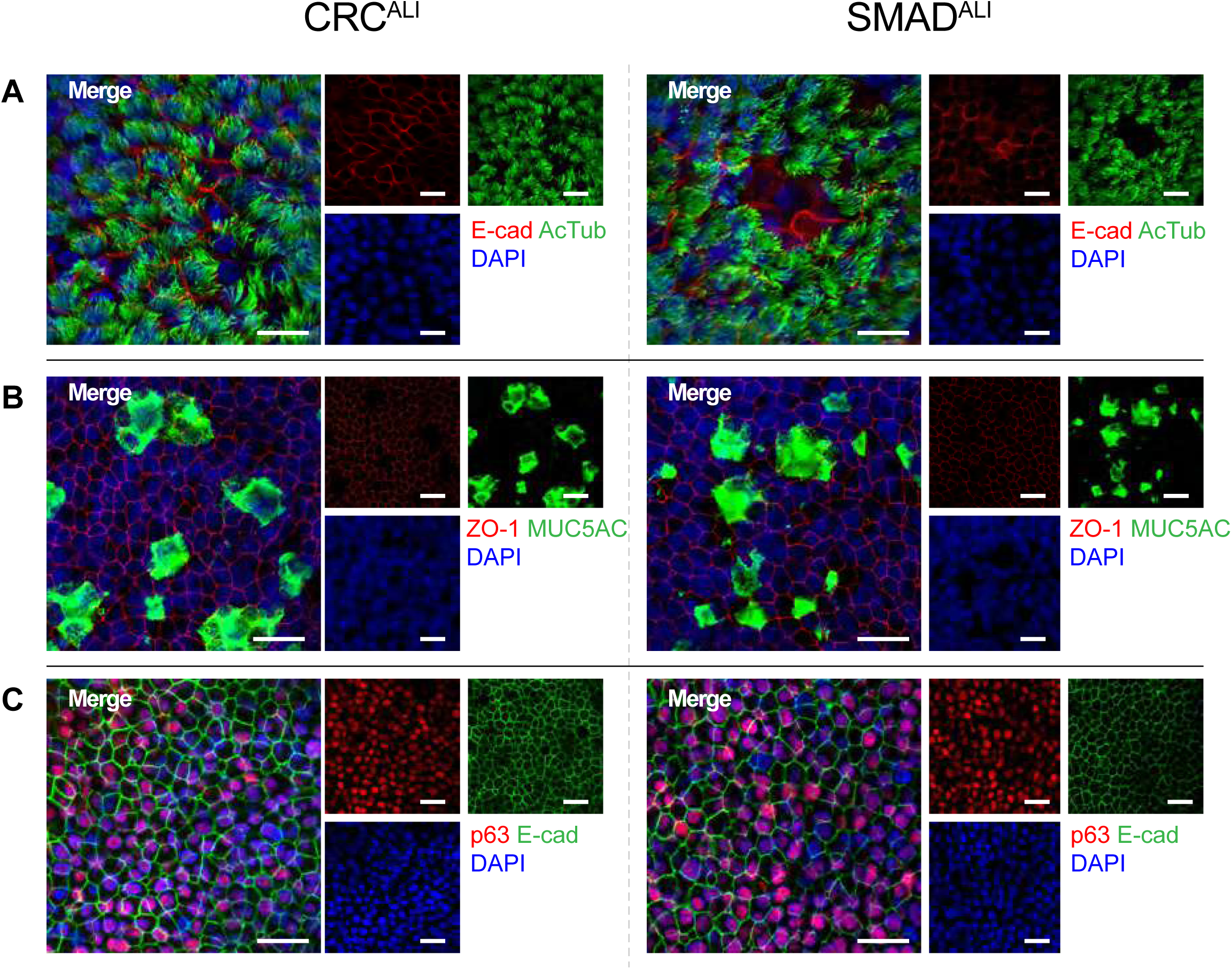
Pseudostratified CRC and SMADi nasal epithelial cells grown at air-liquid interface characteristic of mature airway epithelium. **(A)** Positive staining of acetylated tubulin (ciliated cells; green) and E-cadherin (adherens junction; red) were detected in donor-matched CRC (left panels) and SMADi ALI (right panels) after 21 - 24 days culture. **(B)** Robust expression of MUC5AC (mucus-producing goblet cells; green) and ZO-1 (tight junction; red) were also present. **(C)** p63 (airway basal cell; red) and E-cadherin (green) staining. Images are from Donor 4, F508del/F508del (63x/1.4 oil immersion objective, Zoom 1x). Scale bars = 20µm.

**Fig S2.**
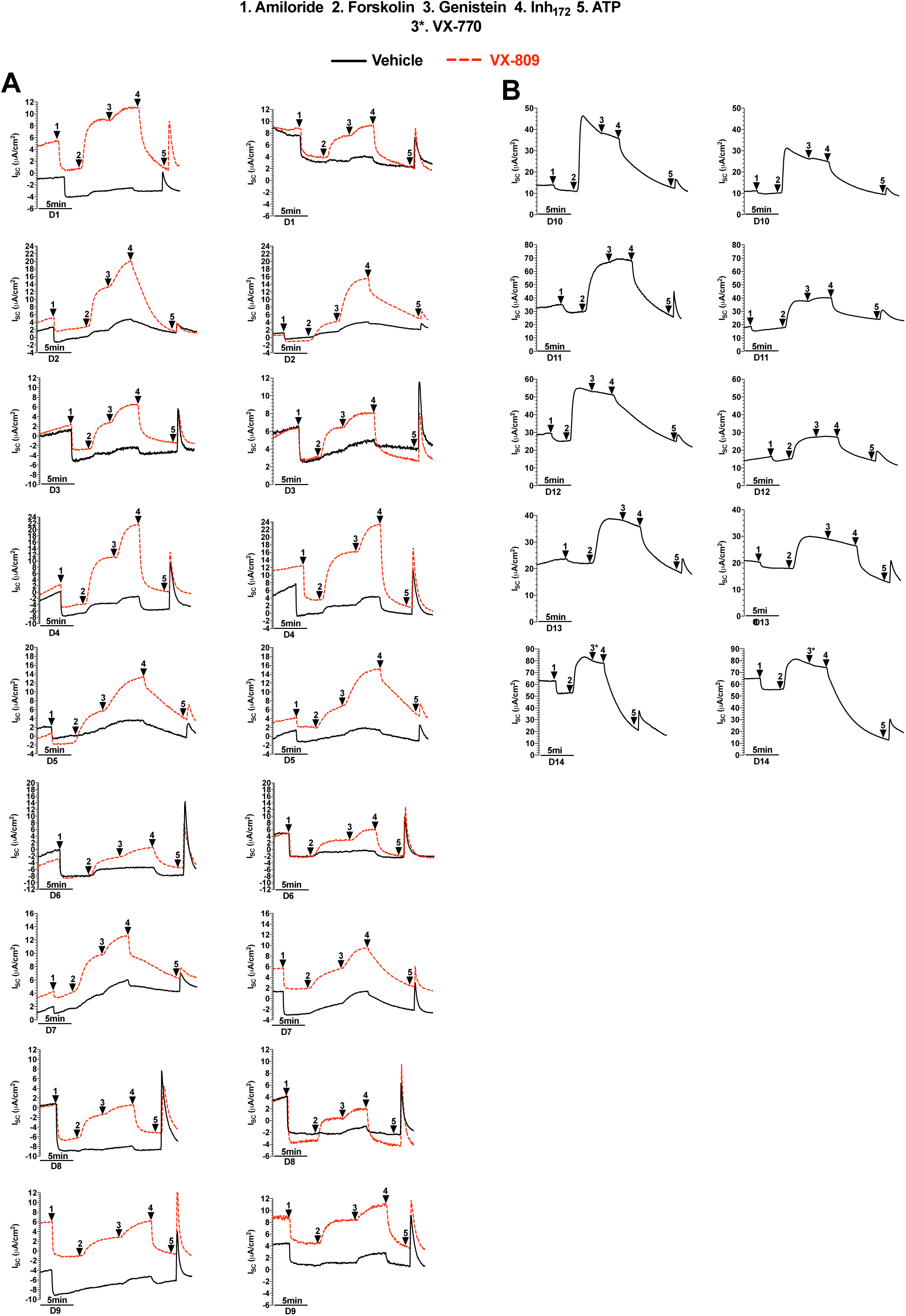
Representative Ussing Chamber short circuit current (Isc) tracings recorded at 37°C for CRC and SMADi expanded HNE ALI cultures. **(A)** CF F508del/F508del-CFTR (D1 to D9). **(B)** Non-CF (wt-CFTR) HNE’s (D10 to D14).

**Fig S3.**
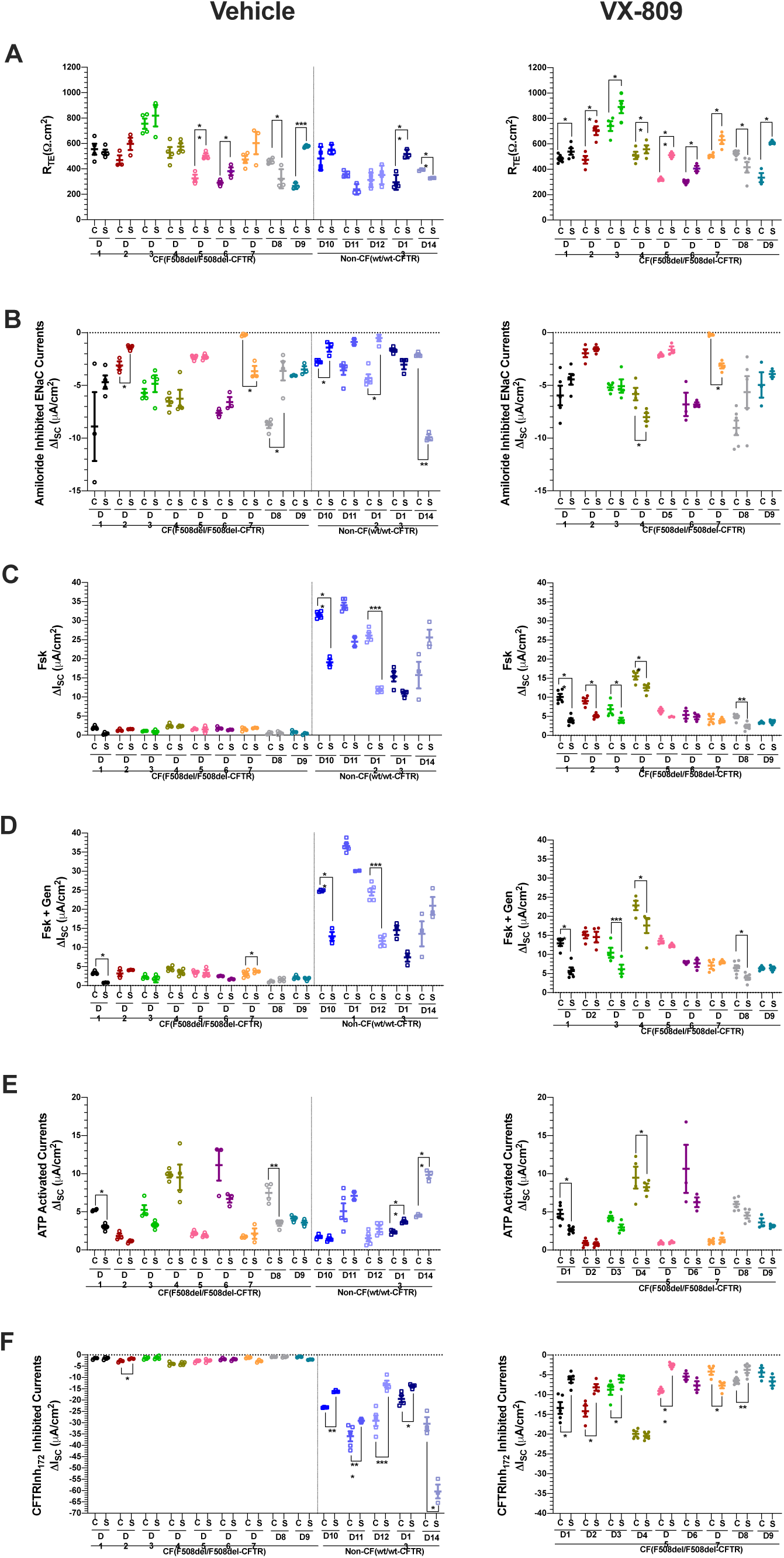
Intra donor comparison of CRCALI vs. SMADiALI cultures. **(A)** Trans-epithelial electrical resistance (RTE), **(B)** Amiloride-inhibited ENaC current, **(C)** Forskolin-stimulated current, **(D)** Forskolin+Genistein-stimulated current, **(E)** ATP-activated currents, **(F)** CFTRInh-172-inhibited current from nine CF donors (D1 to D9) and five non-CF donors (D10 to D14) with or without VX-809 treatment were shown. Each donor is coded with different colour and dotted line separates CF from non-CF donors. Open dots represent vehicle (DMSO) treatment and filled dots represent VX-809 treatment for CF donors. Open squares indicate non-CF donors. Data are presented as means ± standard error of the mean (SEM). Statistical significance is presented as follows: * = P ≤ 0.05; ** = P ≤ 0.01 and *** = P ≤ 0.001.

**Fig S4.**
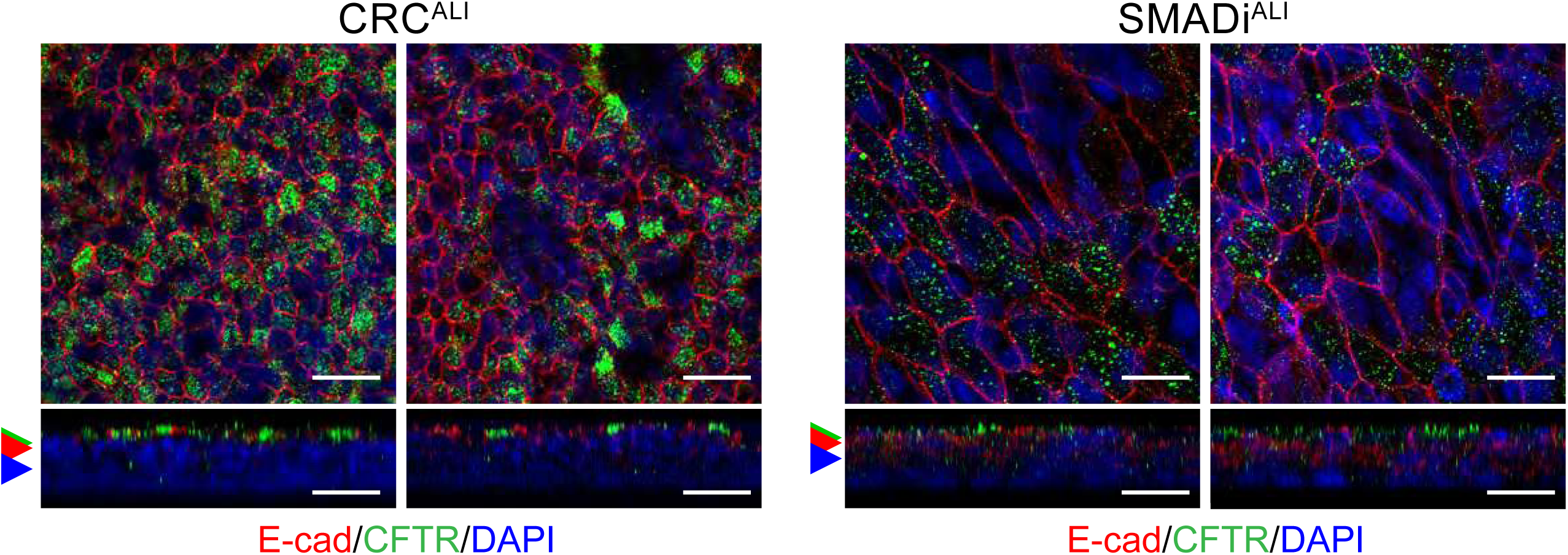
Apical CFTR expression in CRC^ALI^ and SMADi^ALI^ from a non-CF donor (Donor 11). Two different fields of view of CFTR staining are shown for each culture method. XY-images shown in all panels are merged from single channel images acquired at Z-planes indicated by coloured arrows. 63x/1.4 oil immersion objective. Scale bars = 20µm.

